# CRISPR/Cas9-mediated deletion of Shp1 and Shp2 reveals distinct roles in human megakaryopoiesis and proplatelet formation

**DOI:** 10.64898/2026.07.23.735226

**Authors:** Eugénie Schaeffer, Elsa Barré, Desline Hennequin, Cécile Loubière, Lea Mallo, Catherine Strassel, Christian A. Di Buduo, Alessandra Balduini, Yotis A. Senis, Alexandra Mazharian

## Abstract

The non-receptor protein-tyrosine phosphatases Shp1 (*PTPN6*) and Shp2 (*PTPN11*) play critical roles in hematopoietic signaling networks, yet their specific functions in human megakaryopoiesis and thrombopoiesis remain incompletely understood. While Shp2 is recognized in murine models as a positive regulator of thrombopoietin (Tpo)-mediated signaling through the Ras/MAPK and PI3K/AKT pathways, Shp1 has been implicated in RhoA-dependent cytoskeletal remodeling. However, the extent to which these roles translate to human megakaryocyte (MK) development and platelet production is not known. In this study, we systematically investigated the contributions of Shp1 and Shp2 to human MK development and function using CRISPR/Cas9-mediated gene deletion of *PTPN6* and *PTPN11* in CD34^+^ hematopoietic stem and progenitor cells (HSPCs), combined with pharmacological inhibition of Shp2 using the structurally-distinct allosteric inhibitors SHP099 and RMC-4550. Efficient gene editing of *PTPN6* and *PTPN11* resulted in efficient ablation of Shp1 and Shp2 in CD34^+^ HSPC-derived MKs. Genetic deletion or pharmacological inhibition of Shp2 markedly impaired MK proliferation, polyploidization, maturation, and proplatelet formation, whereas loss of Shp1 expression did not. Further, Shp2 inhibition significantly reduced platelet production in a 3-dimensional human bone marrow tissue model. Deletion and inhibition of Shp2 abrogated Tpo-induced ERK1/2 and AKT phosphorylation, confirming its essential role in Mpl receptor signaling. These findings demonstrate the distinct functional roles of Shp1 and Shp2 in MKs and establish Shp2 as a critical positive regulator of Mpl-mediated megakaryopoiesis and thrombopoiesis.

**Key Points:**

- Efficient deletion of Shp1 and Shp2 in human CD34 progenitor cell-derived MKs using CRISPR/Cas9.
- Loss of Shp2 expression impairs thrombopoietin-induced human MK maturation, proplatelet formation and Mpl signaling.

## Introduction

Megakaryopoiesis is the process by which hematopoietic stem and progenitor cells (HSPCs) differentiate into mature megakaryocytes (MKs) that ultimately release platelets into the circulation.^(1,2)^ This process is tightly regulated by cytokines and their respective surface receptors, intracellular signaling pathways and transcriptional regulators, of which thrombopoietin (Tpo) and its receptor Mpl are essential.^(3)^ The three main signaling axes downstream of Mpl are Janus tyrosine kinase/signal transducers and activators of transcription (JAK/STAT), phosphatidylinositol/RAC(RhoA)-alpha serine/threonine protein kinase (PI3K/AKT), and rat sarcoma virus guanosine triphosphatase/mitogen-activated protein kinase (Ras/MAPK), which collectively are critical for MK proliferation, maturation, and proplatelet formation.^(4)^ Despite extensive knowledge of these signaling pathways, the regulatory functions of protein-tyrosine phosphatases (PTPs) in human megakaryopoiesis remain incompletely understood.

The Src homology 2 (SH2) domain-containing non-transmembrane PTPs 1 and 2 (Shp1 and Shp2), encoded by *PTP non-transmembrane 6* and *11* (*PTPN6* and *PTPN11*), respectively modulate signaling from a variety of receptors in hematopoietic cells.^(5)^ Shp1 is a negative regulator of cytokine and immune receptor signaling in myeloid and lymphoid lineages, Shp2 is a positive regulator of cytokine and growth factor receptor signaling in all cells.^(6,7)^ Studies in transgenic mouse models have highlighted essential, but distinct roles of Shp1 and Shp2 in hematopoietic development and function, including megakaryopoiesis and thrombopoiesis.^(8–10)^ However, the specific functions of Shp1 and Shp2 in regulating these processes in human HSPCs and MKs remain undefined.

In this study, we provide the first targeted deletion of *PTPN6* and *PTPN11* in primary human CD34 HSPC-derived MKs using clustered regularly interspaced short palindromic repeats (CRISPR)/CRISPR-associated protein 9 (Cas9) gene editing ^(11)^, enabling a systematic investigation of expression dynamics and functions of Shp1 and Shp2 in human MKs. In parallel, we employ structurally-distinct Shp2-specific allosteric inhibitors to validate and complement the genetic findings.^(12)^ Using various *in vitro* culture conditions, we evaluate the consequences of loss of Shp1 and Shp2 expression on MK maturation, proplatelet formation, platelet production, and Tpo-mediated intracellular signaling. We demonstrate that Shp2, but not Shp1, is indispensable for MK proliferation, endomitosis, proplatelet formation, and platelet release. Shp2 deletion severely impairs Tpo-induced ERK1/2 and AKT phosphorylation, while preserving early lineage commitment and surface receptor expression. Pharmacologic inhibition of Shp2 similarly disrupts terminal MK differentiation, proplatelet formation and platelet production, establishing Shp2 as an essential positive regulator of late-stage human megakaryopoiesis and thrombopoiesis.

## Materials and Methods

### Primary human MK culture

CD34^+^ HSPCs were extracted from leukodepletion filters (TACSI, Terumo BCT, Zaventem, Belgium) and cultured for 14 days, as previously described.^(13,14)^ Cells were seeded at a density of 4 × 10^4^ viable cells/mL in serum-free hematopoietic cell expansion media containing penicillin-streptomycin-glutamine (PSG), 20 µg/mL human low-density lipoprotein (hLDL) and a cytokine cocktail optimized for MK expansion and incubated at 37°C with 5% CO_2_ for 7 days. Cells subsequently matured in serum-free hematopoietic cell expansion media containing PSG, 20 µg/mL hLDL and 50 ng/mL Tpo and incubated at 37°C with 5% CO_2_ for 7 days.

### Reverse transcription-polymerase chain reaction (RT-PCR)

Human cells were collected at days 0, 3, 7, 10, and 12, and washed once in phosphate-buffered saline (PBS) at 4 °C. Cells were subsequently pelleted by centrifugation at 400 × g for 5 min at 4 °C. Total RNA was extracted from freshly isolated CD34 progenitor cells using the RNeasy® Mini Kit (Qiagen). RNA (250 ng) was reverse transcribed with SuperScript III reverse transcriptase (Invitrogen) in a final reaction volume of 30 µL. Quantitative PCR was performed using the RT² SYBR Green qPCR Master Mix (Qiagen) on a CFX96 Touch Real-Time PCR Detection System (Bio-Rad). Relative transcript levels were determined using the 2−ΔΔCt method with *TBP* as the reference gene. Primers for *TBP*, *PTPN6 and PTPN11* were purchased from Genecopoeia. **(Supplemental Table 1)**.

### CRISPR/Cas9 deletion of *PTPN6* and *PTPN11* in CD34^+^ HSPCs

Targeted disruption of *PTPN6* and *PTPN11* was performed in human CD34^+^ HSPCs, using CRISPR/Cas9 technology as previously described.^(15)^ CD34^+^ HSPCs were electroporated at day 2 with ribonucleoprotein complexes consisting of recombinant Cas9 nuclease and sgRNAs targeting *PTPN6* or *PTPN11* using a Lonza 4D-Nucleofector™ **(Supplemental Table 2)**. Cells were cultured in StemSpan™ SFEM II medium (STEMCELL Technologies) supplemented with cytokines. Editing efficiency was assessed by quantifying the protein level by automated Western blotting analysis with antibodies against Shp1 (*PTPN6*) and Shp2 (*PTPN11*) **(Supplemental Table 3)**.

### Ploidy and platelet release

CD34^+^ HSPCs were cultured and differentiated for 14 days. Expression of CD41a and CD42b or CD42c was measured by flow cytometry (BD Fortessa X20 or BD FACSLyric), as previously described.(13,14) MK ploidy was assessed after 10 days of culture by DNA staining with Hoechst dye. Cell viability was determined at day 7 or 10 using FITC Annexin V Apoptosis detection kit (BD Pharmingen). MK proplatelet formation was evaluated at day 14 as previously described.^(16)^ Proplatelet release was induced by manually pipetting cells five consecutive times through a 1 mL pipette tip. Platelets were stained with CD41a (integrin αIIb subunit) Alexa 488- or CD42c (GPIbβ) Alexa 647-conjugated antibodies. (Supplemental Table 4).

### Immunoblotting

Whole cell lysates (WCLs) were prepared from matures MK after 10 days of culture and analyzed by automated capillary-based immunoassay (ProteinSimple Jess), according to manufacturer’s instructions, as previously described.^(17,18)^ Briefly, WCLs were diluted to the required concentration with 0.1X sample buffer, followed by addition of 5X master mix containing 200 mM dithiothreitol (DTT), 5× sample buffer, fluorescent standards and boiled for 5 minutes at 95°C. Samples containing antibody diluent, primary and anti-rabbit secondary antibodies, luminol S-peroxide mix, stripping and wash buffers, were loaded on Jess 12-230 kDa prefilled microplates. Primary antibodies were incubated and the High Dynamic Range (HDR) profile was used for chemiluminescent and fluorescent multiplexing signal detection. Optimized antibody dilutions and sample concentrations used are provided in **(Supplemental Table 4**).

### 3-dimensional (3D) bone marrow tissue model

Megakaryocytes (MKs) were cultured in a porous silk fibroin-based 3D scaffolds to mimic the human bone marrow microenvironment, as previously described.^(20,21)^ Released platelets were analyzed by flow cytometry as described above. Scaffolds were fixed and stained as previously described.^(21)^ Briefly, samples were probed with anti-CD61 (clone SZ21, from Immunotech), and then immersed in Alexa Fluor secondary antibody. For silk fibroin scaffolds imaging, we took advantage of silk auto-fluorescence in the UV light. 3D reconstruction and image processing were performed using Leica licensed software or ImageJ software.

### Statistical analysis

Statistical analysis was performed using GraphPad Prism 10 (GraphPad Software, La Jolla, CA, US). Data was expressed as mean ± standard error mean (SEM). Statistical significance was analyzed by one- or two-way ANOVA followed by the appropriate *post hoc* test, or Kruskal-Wallis test for nonparametric data, a Bonferroni correction was applied for multiple comparisons, as indicated in figure legends. P-values < 0.05 were considered statistically significant.

## Results

### Shp1 and Shp2 expression during human megakaryopoiesis

To establish the expression dynamics of Shp1 and Shp2 during human megakaryopoiesis, we quantified transcript and protein levels of Shp1 and Shp2 in CD34^+^ HSPCs undergoing MK differentiation. Quantitative RT-PCR revealed upregulation of Shp1 transcript at day 3 and return to basal level from day 7 to 10 of differentiation, whereas Shp2 transcript was consistently expressed throughout differentiation **(Figure 1Ai-ii)**. Coincidently, immunoassay revealed that Shp1 protein expression increased during MK differentiation, while SHP2 expression remained constant **(Figure 1Bi-ii)**. These observations indicate that both phosphatases are expressed but differentially regulated during MK development, suggesting unique functions.

**Figure 1.**
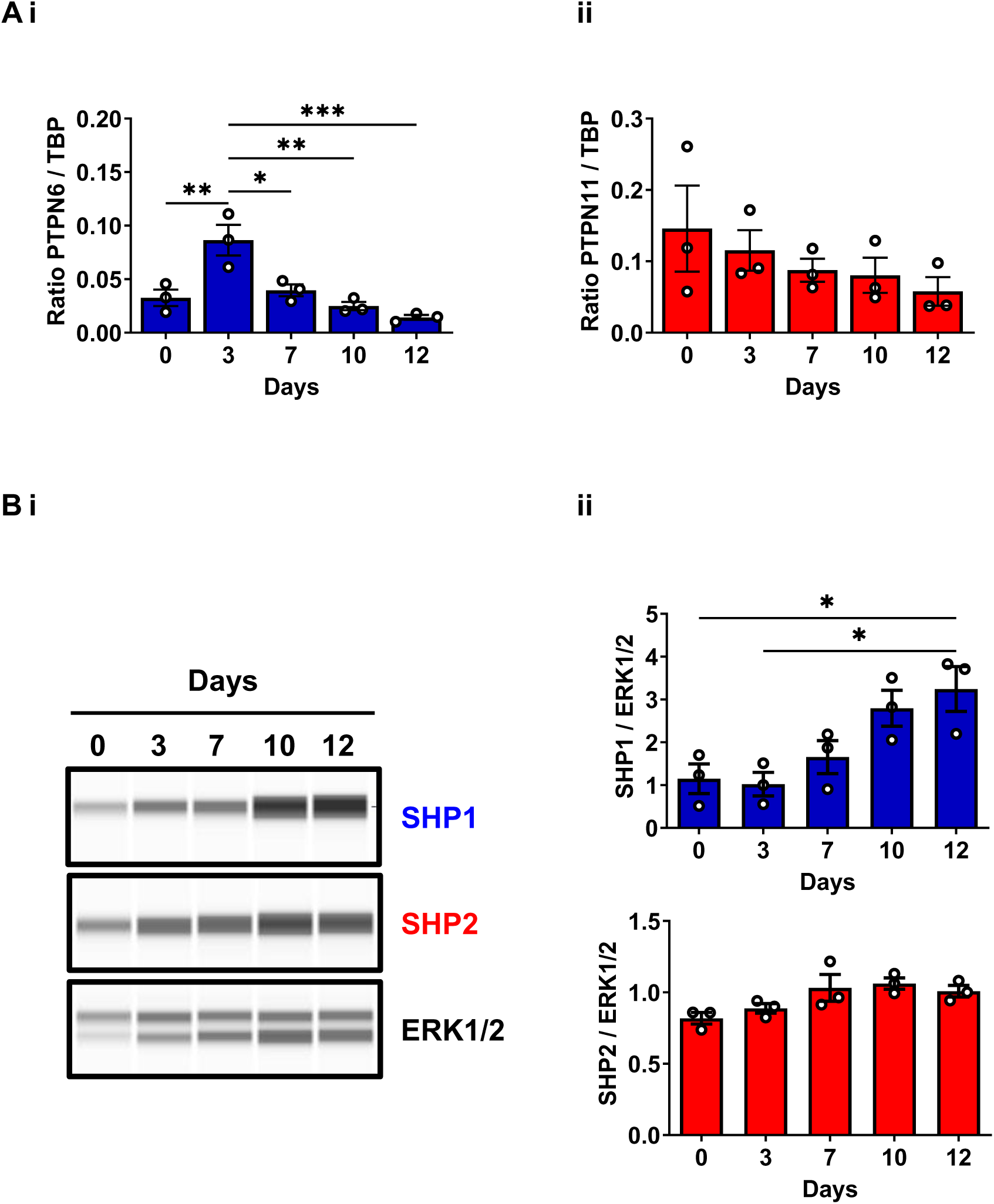
Shp1 and Shp2 expression during human megakaryopoiesis. **(A)** Kinetic expression of *PTPN6* (Shp1) and *PTPN11* (Shp2) genes studied by RT-PCR during megakaryopoiesis. (**B)** Kinetic of Shp1 and Shp2 protein expression during megakaryopoiesis. Representative blots of Shp1 and Shp2 expression levels by capillary immunoassays with the respective antibodies and quantification are shown by immunodetection. n = 3 per condition. Mean ± SEM; one way-ANOVA Kruskal-Wallis; * *P* < 0.05, ** *P* < 0.01, *** *P* < 0.001.

### CRISPR-mediated deletion of Shp1 and Shp2 in human HSPCs

To investigate the function of Shp1 and Shp2 in human megakaryopoiesis, we employed CRISPR/Cas9 gene editing to specifically delete *PTPN6* and *PTPN11* in CD34^+^ HSPCs **(Figure 2A)**. Immunoassay analysis on day 7 of differentiation revealed >90% reduction in Shp1 and Shp2 protein expression compared to control cells, confirming efficient gene ablation **(Figure 2B)**. No compensatory increase in the expression of the non-targeted phosphatase was observed.

**Figure 2.**
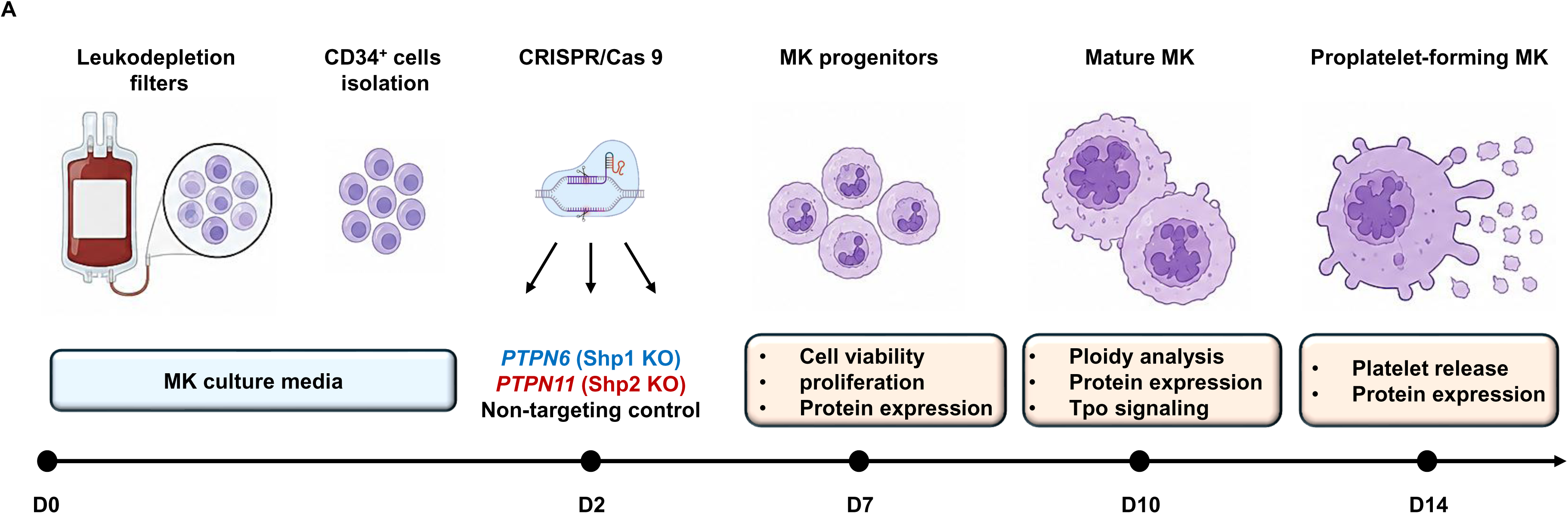
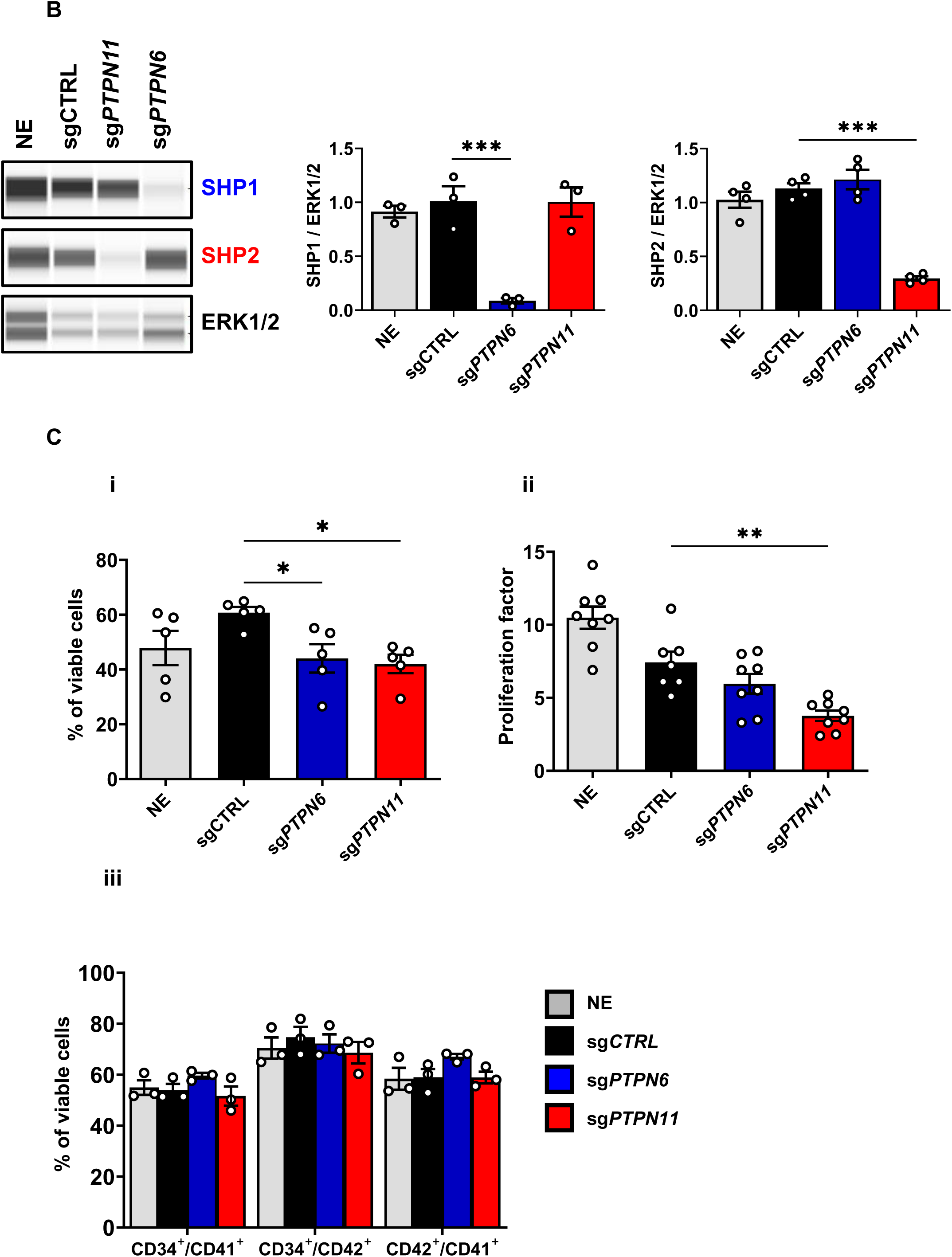
Shp2 KO in CD34^+^ cells reduce megakaryocyte expansion and survival. **(A)** Overview of CRISPR/Cas9 gene editing strategy in CD34^+^ HSPC-derived megakaryocytes (MK). Human CD34^+^-HSPCs were isolated from leukodepletion filters and cultured under megakaryocytic differentiation conditions. On day 3, cells were transfected with ribonucleoprotein (RNP) complexes consisting of Cas9 protein, tracrRNA, and gene-specific crRNA targeting *PTPN6*, *PTPN11*, or a non-targeting control, or non-electroporated (NE). Functional and molecular analyses were performed at sequential stages: cell viability, proliferation, and protein expression on day 7; ploidy, protein expression, and Tpo-induced signaling on day 10; and platelet release and protein expression on day 14. **(B)** Efficient protein knockout following CRISPR/Cas9. Representative blots and densitometry analysis of Shp1 and Shp2 protein expression levels by capillary immunoassays on day 7 MKs, after targeting by negative control or the indicated gene specific CRISPR on day 3. Data are presented as mean ± standard error of the mean (SEM) (3 independent cords per group). 2- way ANOVA to sgCTRL \*\*\**P* < 0.001. (**Ci)** Cell viability was measured with Annexin V and DAPI co-staining. Data are presented as mean ± SEM (3 independent experiment per group). 2-way ANOVA to sgCTRL \**P* < 0.05 **(ii)** Viable cell expansion was assessed by cell count and shown as mean ± SEM (Kruskal-Wallis) \*\**P* < 0.01. **(iii)** Cell phenotype was assessed by measuring CD41a and CD42c surface marker expression by flow cytometry and is shown as mean ± SEM 2-way ANOVA to sgCTRL.

To assess the functional impact of Shp1 and Shp2 deletion on early MK development, we measured cell viability and proliferation following CRISPR/Cas9 gene editing. Deletion of Shp1or Shp2 caused a moderate reduction in viable cell numbers **(Figure 2Ci)**. Importantly, Shp2 deletion markedly reduced cell proliferation, as evidenced by reduced expansion of MK cultures compared to control cells **(Figure 2Cii)**. In addition, surface expression of defining MK receptors, including CD41a and CD42c, was not significantly affected by the deletion of Shp1 or Shp2 **(Figure 2Ciii)**, suggesting that early lineage identity was not affected.

### Shp2 is essential for human MK maturation and proplatelet formation

We also examined the impact of Shp1 and Shp2 deletion on MK development. Flow cytometry analysis at day 10 of differentiation showed that Shp2-deficient MKs exhibited a higher proportion of 2N cells and a sharp reduction in ≥4N cells, indicating a defect in terminal differentiation, whereas Shp1-deficient MKs did not show any detectable differences in ploidy compared to control cells **(Figure 3A)**. Immunoassay analysis confirmed that Shp1 and Shp2 protein levels remained >90% depleted at this stage of development **(Figure 3B)**. Further, there is a significant reduction in the number of platelets released by Shp2-, but not Shp1-deficient MKs, as assessed by flow cytometry at day 14 **(Figure 3Ci)**. Shp1 and Shp2 protein levels remained >90% depleted at this time point **(Figure 3Cii)**, demonstrating sustained gene knockout and supporting a direct role for Shp2 in regulating proplatelet formation at this late stage of differentiation.

**Figure 3.**
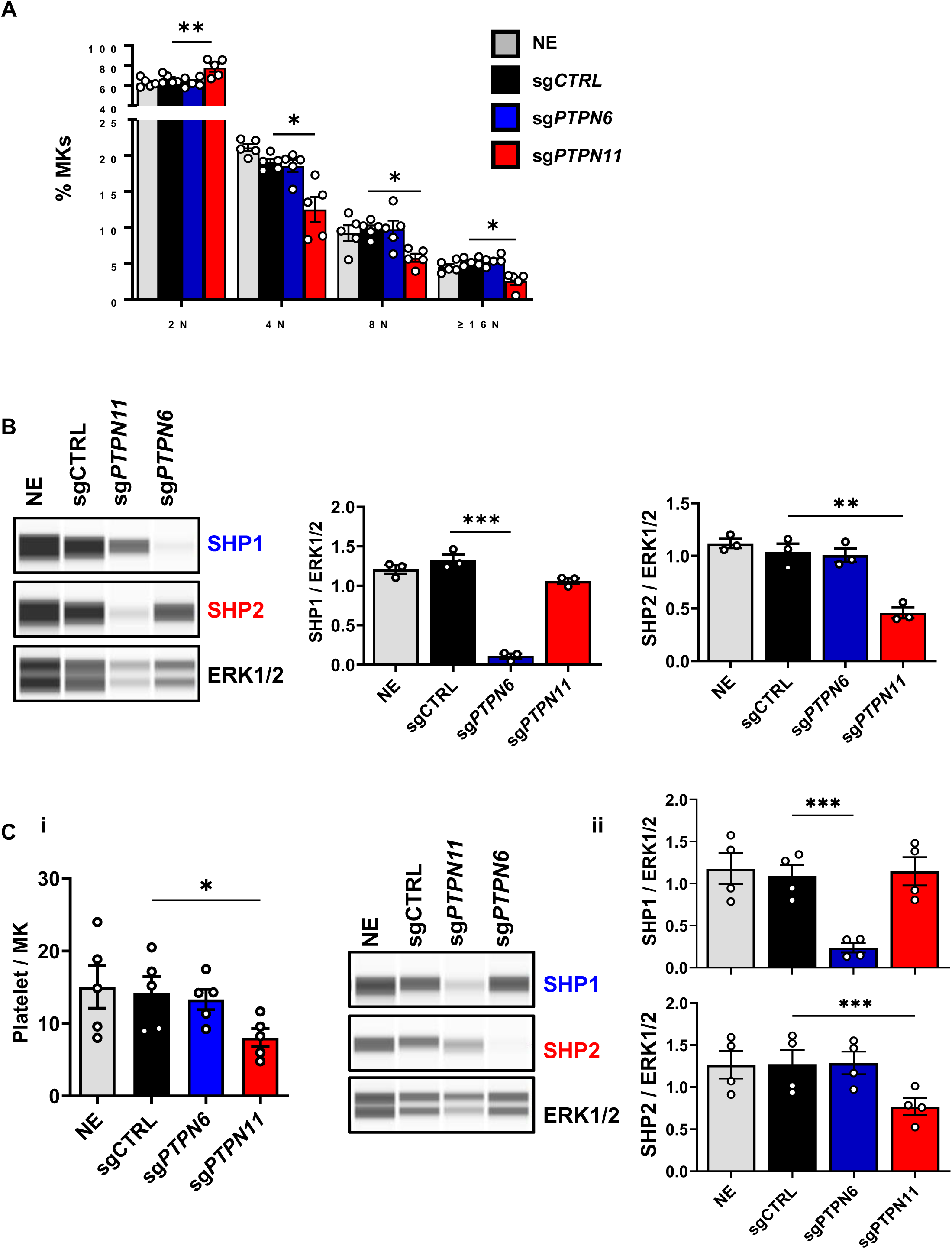
Shp2 deletion impairs megakaryocyte maturation and platelet release. **(A)** Cell ploidy was measured by flow cytometry using Hoechst and propidium iodide co-staining and is shown as mean ± SEM 2-way ANOVA to sgCTRL. * p<0.05, **p<0.01. **(B)** Efficient protein knockout following CRISPR/Cas9 at day 10. Representative blots and densitometry analysis of Shp1 and Shp2 protein expression levels by capillary immunoassays on day 10 MKs after targeting by negative control or the indicated gene specific CRISPR on day 3. Data are presented as mean ± SEM (3 independent cords per group). 2-way ANOVA to sgCTRL, ***p<0.001. **(C)** Shp2 deletion impairs platelet release. **(i)** Platelet release was measured by flow-cytometry and is shown with mean ± SEM 2-way ANOVA to sgCTRL. * p<0.05. **(ii)** Efficient protein knockout following CRISPR/Cas9 is maintained at day 14. Representative blots and densitometry analysis of Shp1 and Shp2 protein expression levels by capillary immunoassays on day 14 MKs after targeting by negative control or the indicated gene specific CRISPR, ***p<0.001.

Collectively, these findings indicate that Shp2 is required for optimal cell proliferation, endomitosis and platelet release during human MK differentiation, while Shp1 is not involved in these early developmental processes under steady-state conditions.

### Deletion of Shp2 impairs Tpo signaling in human MKs

We next analyzed Tpo signaling in Shp1- and Shp2-deficient human MK, at day 10 of culture by capillary-based immunoassay. Following Tpo stimulation, control CD34^+^ HSPC-derived MKs transfected with a non-targeting sgRNA exhibited robust phosphorylation of ERK1/2, AKT, and STAT3, indicating activation of canonical Mpl signaling and consistent with known signaling pathways essential for megakaryopoiesis **(Figure 4A-B)**. In Shp2-deficient MKs, Tpo-induced ERK1/2 phosphorylation was almost completely abrogated under the same conditions, and AKT activation was significantly reduced, demonstrating that Shp2 is a critical regulator of Mpl signaling. Total levels of ERK1/2 and AKT expression remained unchanged, confirming that the observed effects were due to impaired phosphorylation rather than reduced protein expression **(Figure 4A-B)**. In contrast, MKs lacking Shp1 displayed normal phosphorylation of ERK1/2, AKT, and STAT3 in response to Tpo, demonstrating that Shp1 is dispensable for these signaling pathways **(Figure 4A-B)**.

**Figure 4.**
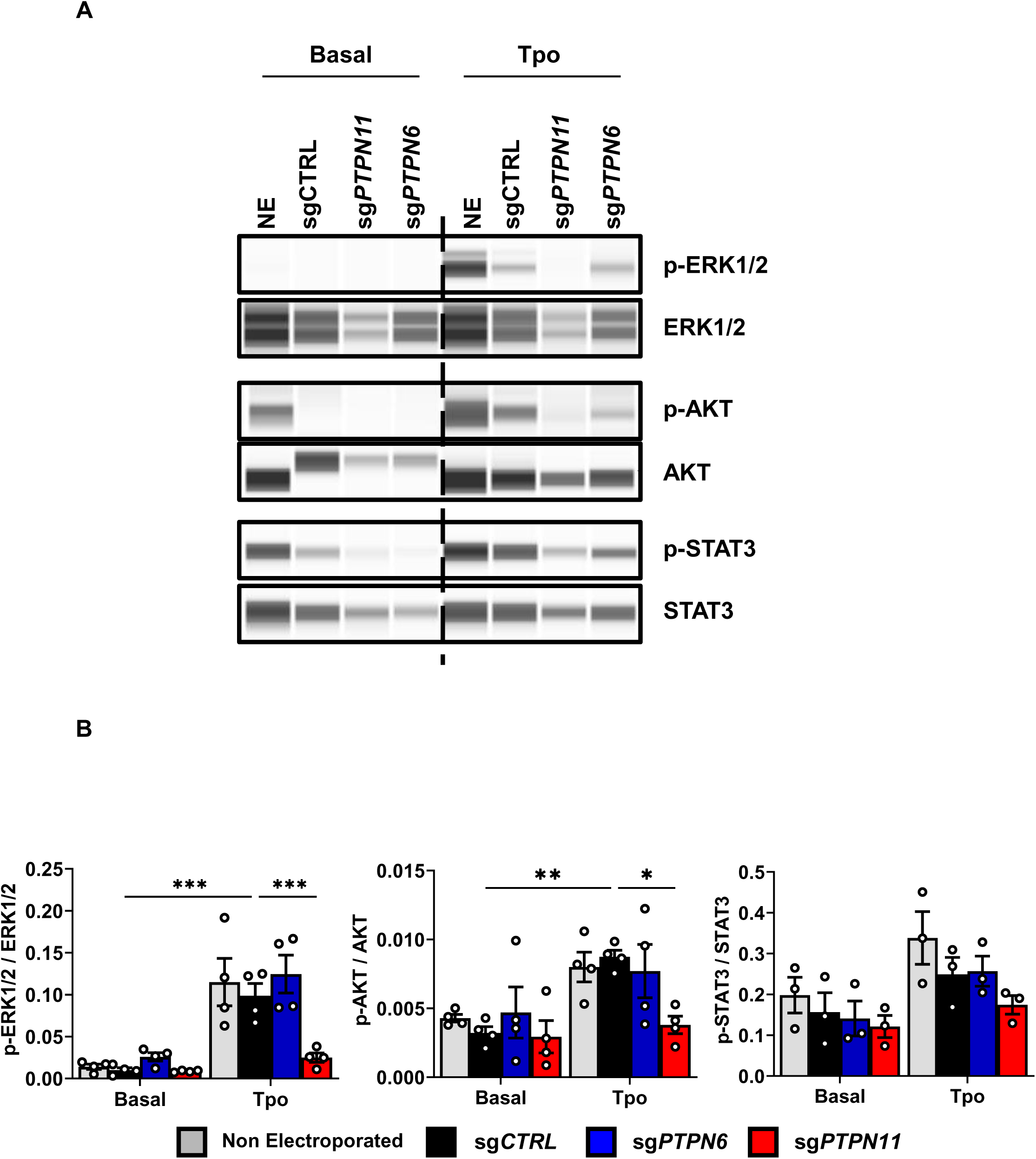
CRISPR-mediated Shp2 KO impairs Mpl signaling in human megakaryocytes. **(A)** Immunodetection of Shp1, Shp2, p-ERK1/2 (T202/Y204), ERK1/2, p-AKT (S473), AKT, p-STAT3 (Y705) and STAT3 following Tpo treatment. **(B)** Phosphorylation of ERK1/2, AKT and STAT3 were quantified and are shown normalized to total protein quantification with mean ± SEM (2-way ANOVA to sgCTRL). **p<0.01, ***p<0.001.

### Shp2 inhibition replicates Shp2 deletion

To validate the genetic findings and further characterize the stage-specific requirement of Shp2 during megakaryopoiesis, we treated CD34^+^ HSPC-derived MKs with the two structurally-distinct Shp2-specific allosteric inhibitors SHP099 and RMC-4550.^(22,23)^ Both compounds selectively stabilize Shp2 in its inactive conformation, preventing it from dephosphorylating downstream phosphoproteins. Both inhibitors significantly reduced cell proliferation, consistent with the role of Shp2 in promoting cytokine-driven cell cycle progression **(Figure 5Ai-ii)**. Flow cytometry analysis revealed a block in endomitosis, with a reduction in the proportion of ≥8N MKs **(Figure 5Aiii)**. Moreover, proplatelet formation was markedly reduced at day 14 with either inhibitor, as shown by the significant reduction in the number of platelets released by MKs **(Figure 5Aiv)**. Importantly, pharmacologic inhibition of Shp2 also disrupted Tpo-mediated signaling. Human CD34^+^ HSPC-derived MKs treated with SHP099 or RMC-4550 showed a major reduction in Tpo-induced ERK1/2 and AKT phosphorylation **(Figure 5Bi-ii)**, similar to the signaling defects observed in Shp2-deficient MKs and confirming that acute pharmacologic inhibition of Shp2 impairs Mpl signaling and downstream functional responses. Collectively, findings demonstrate that pharmacologic inhibition of Shp2 reproductions key aspects of the Shp2-deficient phenotype, further validating the non-redundant and stage-specific role of Shp2 in human MK development, as well as the importance of its catalytic activity. It also suggests that Shp2 expression alone is insufficient to sustain its functions.

**Figure 5.**
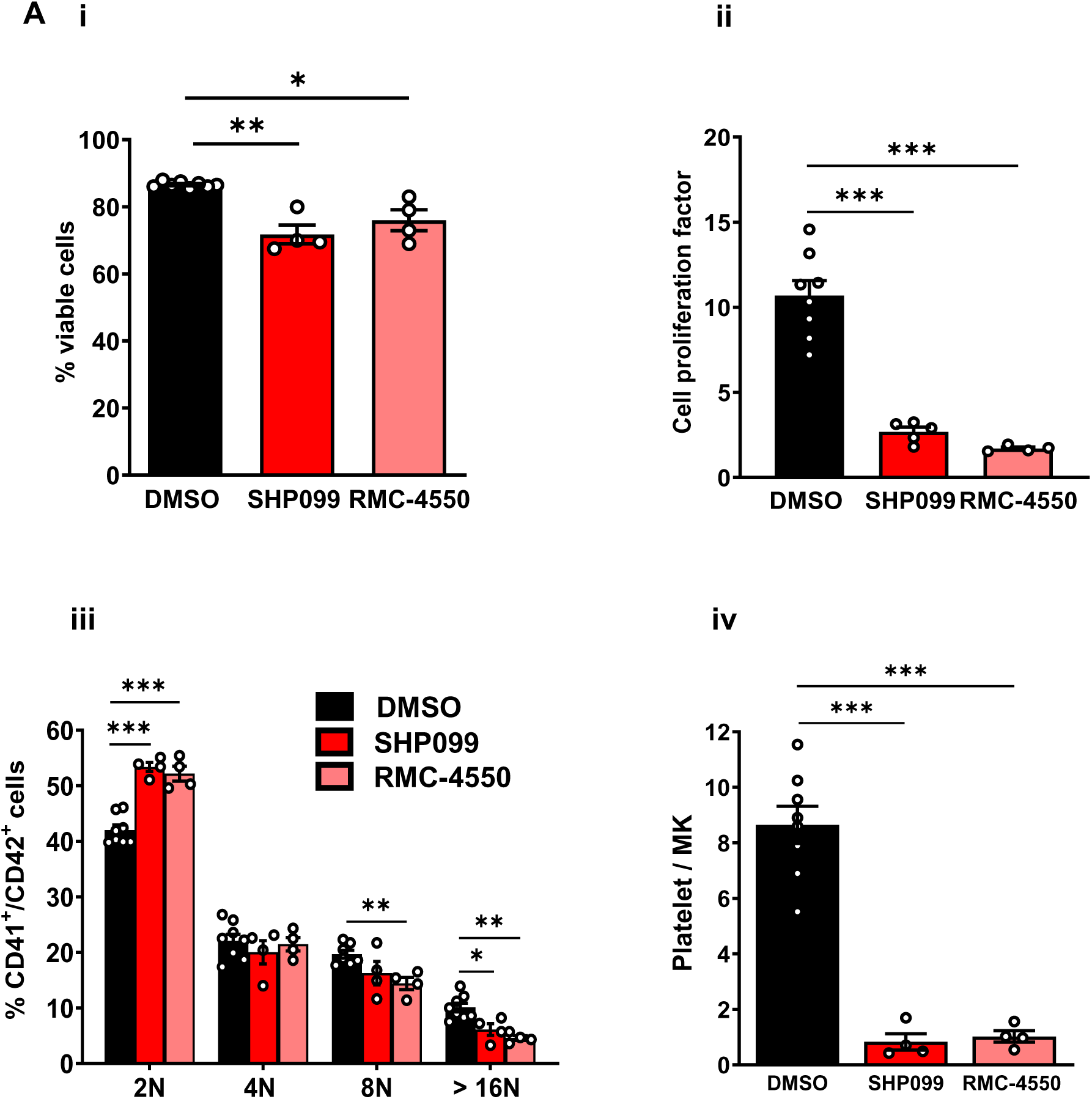
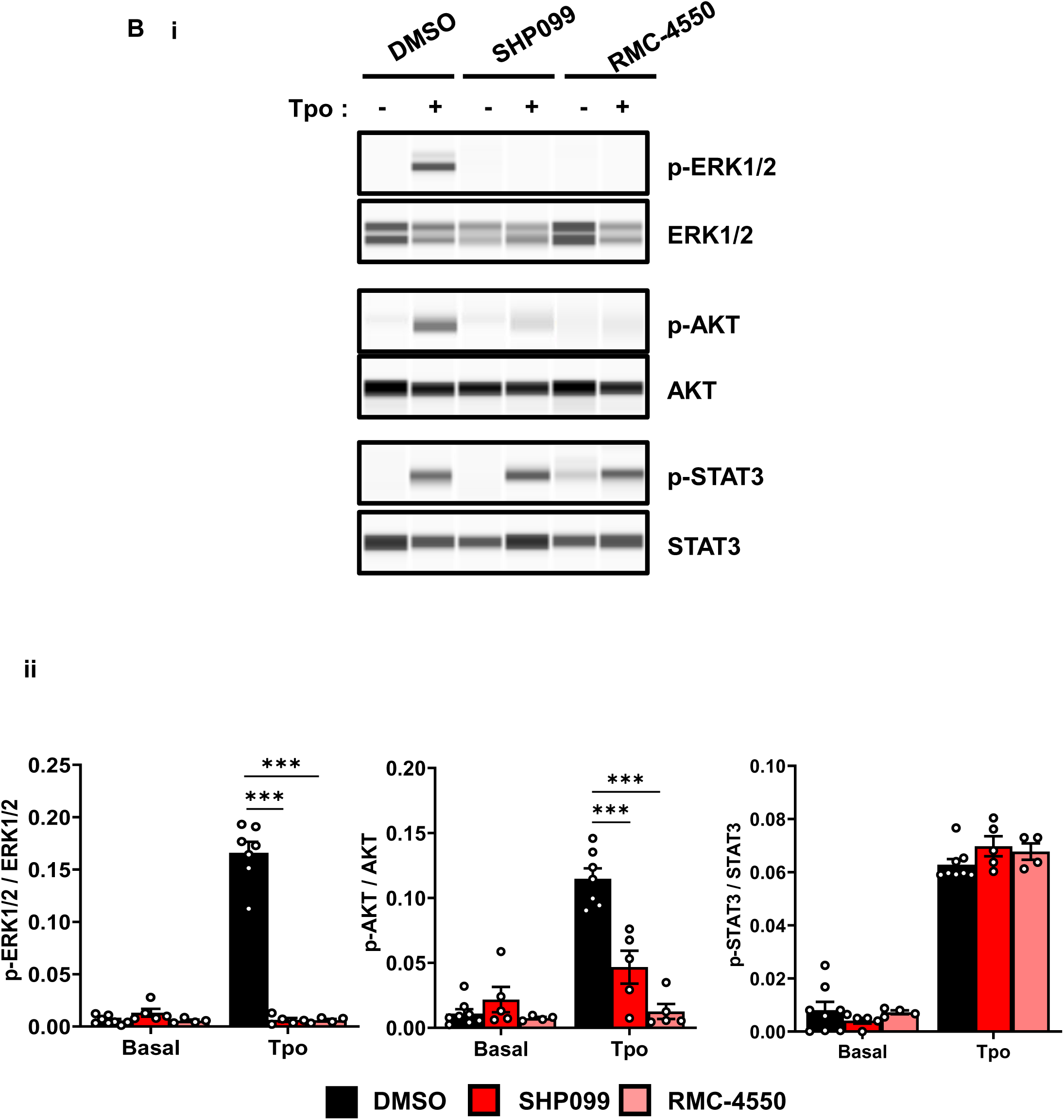
Pharmacological inhibition of Shp2 impairs human MK maturation and Mpl signaling. **(A) (i)** Viable cell expansion was assessed by cell count and shown with mean ± SEM (Unpaired t-test). **(ii)** Cell proliferation was measured assessed by cell count and is shown with mean ± SEM (Unpaired t-test). **(iii)** Cell ploidy was measured by flow cytometry using Hoechst and propidium iodide co-staining and is shown with mean ± SEM (2-way ANOVA DMSO). **p<0.01, ***p<0.001. **(iv)** Proplatelet formation. Proplatelet release was measured by flow-cytometry and is shown with mean ± SEM (Unpaired t-test), ***p<0.001. (**B**) Shp2 inhibitors impairs Mpl signaling. **(i)** Immunodetection of p-ERK1/2 (T202/Y204), ERK1/2, p-AKT (S473), AKT, p-STAT3 (Y705) and STAT3 following Tpo treatment. **(ii)** Phosphorylation of ERK1/2, AKT and STAT3 were quantified and are shown normalized to total protein quantification with mean ± SEM (2-way ANOVA DMSO), ***p<0.001.

### Shp2 is required for platelet production in a 3D bone marrow model

To evaluate the physiological relevance of our findings, we next examined Shp2 inhibition across progressive stages of thrombopoiesis using both 2D and 3D models, as previously described.^(24,25)^ Pharmacological inhibition of Shp2 in 2D liquid cultures with either SHP099 or RMC-4550 resulted in defective proplatelet formation by day 14 of differentiation **(Figure 6A)**. To further explore terminal MK function, CD34^+^ HSPC-derived MKs were cultured on fibronectin-coated surfaces, a model that promotes cytoskeletal re-organization and proplatelet bifurcation. Under these conditions, Shp2 inhibition resulted in a significant reduction in the proportion of proplatelets, bifurcations and elongation of cytoplasmic extensions, hallmarks of pre-platelet and proplatelet maturation **(Figure 6B)**. These morphological defects were consistent with both allosteric inhibitors (data not shown). To assess platelet release in a simulated bone marrow microenvironment, a perfusable 3D silk fibroin-based model was employed.^(26)^ MKs treated with either SHP099 or RMC-4550 were seeded in the silk scaffolds and cultured under flow. Quantitation of platelet-sized CD41^+^CD42b^+^ particles released into the circulation channels revealed a significant reduction in platelets released by Shp2-inhibited cultures compared to untreated controls **(Figure 6C)**.

**Figure 6.**
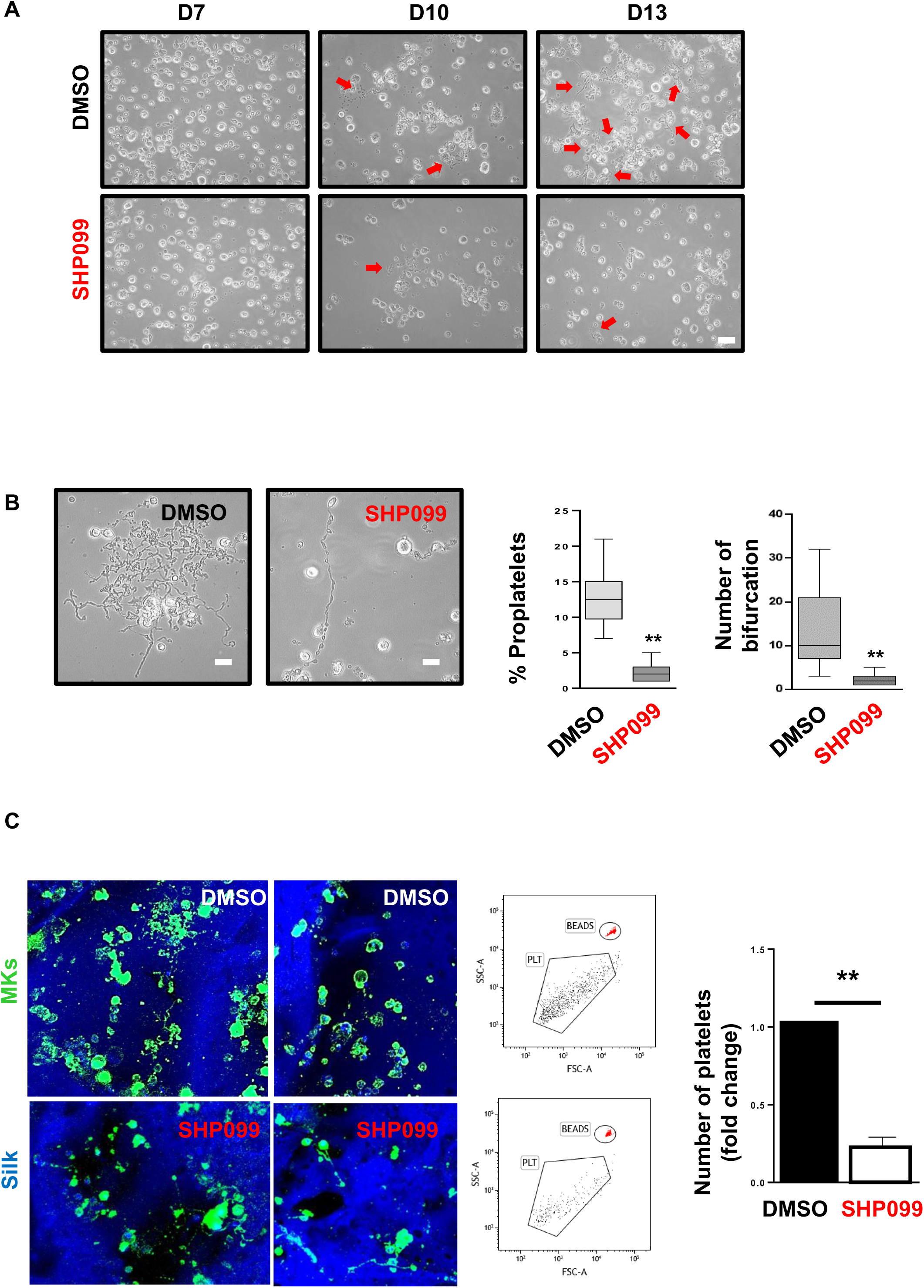
SHP099 inhibits proplatelet formation and platelet release from megakaryocytes in a 3D silk bone marrow model. **(A)** Representative bright-field images of CD34^+^-derived MKs treated with DMSO or SHP099. Red arrows indicate proplatelet extensions. Scale bar: 50 μm. **(B)** Quantification of proplatelet-forming MKs per field. Data are shown as mean ± SD from n = 3 independent experiments. P < 0.01, unpaired t-test. Scale bar: 25 μm. **(C)** Immunofluorescence images of MKs embedded in silk-based 3D scaffolds treated with DMSO or SHP099. MKs are labeled in green (CD61); nuclei are counterstained with DAPI (blue). Scale bar: 100 μm. Flow cytometry analysis of platelets released from silk scaffolds. Platelets (PLT) were gated based on forward and side scatter and quantified using counting beads. SHP099 treatment significantly reduced platelet release compared with DMSO. Data represent n = 3 independent experiments.

These findings confirm that Shp2 catalytic activity is essential for MK maturation and proplatelet formation under different culture conditions, highlighting its central role in late stage megakaryopoiesis, terminal platelet release and Mpl signaling.

## Discussion

In this study, we demonstrate that Shp2 plays a non-redundant and essential role in human MK maturation, thrombopoiesis and Tpo-mediated Mpl signaling. Using both CRISPR/Cas9- mediated gene editing and pharmacologic inhibition in CD34^+^ HSPC-derived MKs, we show that Shp2 loss impairs cell proliferation, endomitosis, proplatelet formation, and platelet production, both in standard 2D cultures and in a physiologically relevant 3D bone marrow model. In contrast, Shp1 appears to be dispensable for MK development under steady-state conditions, highlighting a functional divergence between these structurally related tyrosine phosphatases. Importantly, this study represents one of the first functional applications of CRISPR/Cas9 technology in the context of terminal MK differentiation and platelet production in primary human cells, establishing a valuable platform to interrogate gene function in a lineage that is otherwise difficult to genetically manipulate.

While the roles of Shp1 and Shp2 in murine megakaryopoiesis and thrombopoiesis have been previously reported, the extent to which these findings apply to human MKs has remained unexplored.^(8–10)^ Human MKs differ from their murine counterparts in multiple aspects, including size, ploidy, signaling kinetics, transcriptional programs, and responses to cytokines such as Tpo.^(28,29)^ By performing gene deletion directly in CD34^+^ progenitors prior to differentiation, circumvents these issues, particularly pertaining to ploidy, and provides definitive functional evidence that Shp2 is a non-redundant and stage-specific regulator of human MK development and platelet production. Notably, we show that Shp2 is indispensable for fundamental aspects of megakaryopoiesis, including proliferation, endomitosis, proplatelet formation and platelet release, irrespective of the culture conditions, thereby reinforcing its central role in thrombopoiesis.

Mechanistically, loss of Shp2 expression prevented ERK1/2 phosphorylation and significantly reduced AKT phosphorylation following Tpo stimulation, establishing that it is an essential positive regulator of Mpl signaling.^(30)^ ERK1/2 and AKT expression remained unchanged, ruling out non-specific effects on protein expression and affirming a specific signaling defect.

A notable strength of this study lies in the parallel use of the structurally-distinct Shp2- specific allosteric inhibitors SHP099 and RMC-4550, which not only validate the genetic findings, but also extend their relevance to therapeutic contexts. These inhibitors reproduced the Shp2-deficient phenotype, both functionally and biochemically, reinforcing that Shp2 enzymatic activity is critical for MK maturation, and protein expression is insufficient. As Shp2 inhibitors are currently under clinical development for cancer treatment,^(31,32)^ our findings raise the possibility of inhibiting Shp2 in instances of thrombocytosis, but also concern of inducing thrombocytopenia in patients with normal platelet count. The bleeding risk is low, nonetheless, as evidenced from Shp2 conditional knockout mouse models.^(33)^ Interestingly, Shp1 deletion had minimal effects on MK proliferation, maturation and Tpo signaling, despite its structural similarities with Shp2 and increased expression during early-stage MK differentiation. These findings contrast with murine models where Shp1 has been implicated in the negative regulation of immune and hematopoietic receptor signaling pathways, including cytoskeletal reorganization in MKs.^(33)^ Whether Shp1 functions under specific stress or inflammatory contexts in human megakaryopoiesis and thrombopoiesis remains to be determined.

Our study demonstrates the technical feasibility of using CRISPR/Cas9 gene editing in primary human HSPCs to study late-stage MK lineage differentiation, which remains a major challenge in the field. Most functional genomic studies rely on immortalized cell lines or murine models, which can fail to recapitulate specific physiological features of MKs. Our approach allows for high-efficiency gene ablation and robust functional readouts in mature MKs, enabling detailed dissection of gene function in the final stages of thrombopoiesis. This strategy may serve as a powerful approach for studying other regulators of human megakaryopoiesis, congenital thrombocytopenia, or to model disease-associated mutations.

In conclusion, our findings establish Shp2 as a critical and non-redundant effector of human megakaryopoiesis and thrombopoiesis, with a mechanistic link to Tpo/Mpl signaling. These discoveries provide important insight into the lineage-specific function of Shp2 in human hematopoiesis and highlight the translational impact of targeting Shp2 in clinical settings. Future research should examine the role of Shp2 in disease states characterized by altered megakaryopoiesis, such as myeloproliferative neoplasms,^(34,35)^ or congenital thrombocytopenias to assess whether Shp2 modulation could be reversible or selectively targeted without compromising platelet production. More broadly, we introduce a robust CRISPR/Cas9-based system for gene interrogation in human MK that opens new avenues for functional genomics in primary hematopoietic lineages.

## Acknowledgements

This study was supported by Inserm, the Agence Nationale pour la Recherche (AM, ANR-22- REMYS-CE14-0061-01 and YAS, ANR-23-TARGITT-CE18-0020-01), the University of Strasbourg (IdEx 2023 Attractivity) and the European Research Council (YAS ERC- MENTOR-101141783).

## Author Contributions

E.S. Performed experiments, analyzed data, revised the manuscript

E.B. Performed experiments, analyzed data, revised the manuscript

D.H. Performed experiments and analyzed data

C.L. Performed experiments and analyzed data

L.M. Provided technical support

C.S. Provided technical support

C.A.D.B. Performed experiments, analyzed data and revised the manuscript

A.B. Revised the manuscript.

Y.A.S. Designed experiments, analyzed data, wrote and revised the manuscript

A.M. Conceptualized, designed experiments, analyzed data, wrote and revised the manuscript

## Data Availability Statement

The data that support the findings of this study are available from the corresponding author upon reasonable request.

