## Supplemental Figures for "CRISPR/Cas9-mediated deletion of Shp1 and Shp2 reveals distinct roles in human megakaryopoiesis and proplatelet formation"

**Table 1. PTPN6 and PTPN11 primers for qRT-PCR.**

| Gene | Sequence | Forward (5'-3') | Reverse (3'-5') | Reference Biorad |
|---|---|---|---|---|
| **PTPN6** | 5’**TGGTTTCACCGAGACCTCAG**3’ | 5′-TTGACCACAGCCGAGTGATCCT-3′ | 5′-CTGGCGATGTAGGTCTTAGCGT-3′ | ***qHsaCED0043391*** |
| **PTPN11** | 5’**GCGCACTGGTGATGACAAAG**3’ | 5′-TCAGCACAGAAATAGATGTG-3′ | 5′-TGCTTATCAAAAGGTAGTCA-3′ | ***qHsaCID0022662*** |

**Table 2. PTPN6 and PTPN11 single guide RNAs for CRISPR/Cas 9 gene editing.**

| Gene | Synthego | IDT |
|---|---|---|
| **PTPN6** | ***sgPTPN6.1*** PTPN6+6951627 | ***sgPTPN6.3*** Hs.Cas9.PTPN6.1.AE |
| | ***sgPTPN6.2*** PTPN6-6951610 | |
| **PTPN11** | ***sgPTPN11.1*** PTNPN11+112446394 | ***sgPTPN11.2*** Hs.Cas9.PTPN11.1.AA |

**Table 3. Antibodies and working concentrations for JESS experiments.**

| Antibody | Dilution | Protein concentration | References |
|---|---|---|---|
| Rabbit anti-Shp1 | 1/10 | 0.1 mg/ml | 3759<br>Cell Signaling |
| Mouse anti-Shp2 | 1/20 | 0.1 mg/ml | 20145-1-AP<br>ProteinTech |
| p-ERK1/2<br>(Thr202/Tyr204) | 1/25 | 0.1 mg/ml | 9101<br>Cell Signaling |
| ERK1/2 | 1/25 | 0.1 mg/ml | 9102<br>Cell Signaling |
| p-AKT (Ser473) | 1/20 | 0.1 mg/ml | 9271<br>Cell Signaling |
| AKT | 1/50 | 0.1 mg/ml | 4691<br>Cell Signaling |
| p-STAT3 (Tyr705) | 1/10 | 0.1 mg/ml | 9145<br>Cell Signaling |
| STAT3 | 1/50 | 0.1 mg/ml | 4904<br>Cell Signaling |

**Table 4. Antibodies used for flow cytometry.**

| Antibody | Dilution | References |
|---|---|---|
| CD34-PeCy7 | 1/100 | 15598386<br>Invitrogen |
| CD41a-Alexa568 | 1/100 | Invitrogen |
| CD41-APC | 1/100 | B16894<br>Beckman Coulter |
| CD42b-PE | 1/100 | IM1417U<br>Beckman Coulter |
| CD42c-Alexa647 | 1/200 | 25-0349-42<br>Invitrogen |
| AnnexinV-FITC | 1/100 | 51-65874X<br>BD Pharmigen |
